# The Effect of SARS-COV-2 Protein Fragments on the Dimerization of α-Synuclein

**DOI:** 10.1101/2025.08.26.672456

**Authors:** Lucy M Coleman, Ulrich H. E. Hansmann

## Abstract

There is evidence that amyloidogenic segments in SARS-COV-2 proteins can induce aggregation of α-synuclein (αS), the main component of brain-located amyloids whose presence is connected with Parkinson’s Disease (PD). Using molecular dynamic simulations, we could show in earlier work that SARS-COV-2 protein fragments shift the ensemble of αS chains toward more aggregation-prone conformations. However, the mechanism by which these chains assemble into fibrils, the presumed neurotoxic agent in PD, is not clear. The first step on that route are dimers. For this reason, we have now, using again molecular dynamics simulations, studied how the fragment _194_FKNIDGYFKI_203_ (FI10) of the SARS-COV-2 spike protein, and the fragment _54_SFYVYSRVK_62_ (SK9) of the envelope protein, alter the ensemble of α-synuclein dimers. Our simulations suggest a differential stabilization of such dimers that would preferentially seed rod-like fibrils over the competing twister-like structures.

## I. INTRODUCTION

A hallmark of Parkinson’s Disease (PD) is the presence of amyloids located in the brain of patients that are made mainly of α-synuclein (αS) and appear to be the neurotoxic agent.^1, 2^ As there have been correlations observed between falling ill with COVID-19 and outbreaks of PD,^3^ and SARS-COV-2 induced αS amyloid formation has been found *in vitro,*^4^ we and other groups^5^ have proposed that amyloidogenic SARS-COV-2 protein regions can enhance aggregation of αS potentially causing PD. We have speculated that during acute inflammation, as commonly seen in COVID-19, neutrophils release enzymes that cleave SARS-COV-2 proteins into amyloidogenic fragments which in turn cross-seed human proteins. Such cleavage has been shown for the amyloidogenic segment of residues _194_FKNIDGYFKI_203_ (FI10) of the spike protein.^6^ Using all-atom molecular dynamics simulations we have shown that this peptide and the segment _54_SFYVYSRVK_62_ (SK9) of the envelope protein shift the ensemble of αS monomers toward conformations that are more aggregation prone.^7, 8^ However, the likely neurotoxic agents in PD are nor these monomers but their assemblies into fibrils that are characterized by a cross-beta structure; and the pathogenesis and severity of PD is correlated with the structures of the polymorphic fibrils in the patient brains.

The first step on the road to such fibrils are dimers that may serve as their seeds. This is the reason why we investigate in the present study the effect of SARS-COV-2 protein fragment on the stability of the dimers. For this purpose we employ all-atom molecular dynamics simulations where the above two SARS-COV-2 protein fragments interact with αS dimers build from aggregation-prone monomer configurations collected in our earlier work. We are especially interested to see if any stabilizing effect would depend also on the αS dimer structure, and therefore also leading to specific fibril forms, or if the earlier observed differential stabilization of fibril polymorphs^8^ depends only on the energetics of the fibril conformations but not on the kinetics of their formation.

## II. RESULTS AND DISCUSSION

In previous work we could show that interaction with SARS-COV-2 protein fragments such as SK9 or FI10 shift the ensemble of αS chains toward more extended and aggregation prone conformations. We have argued that the observed distribution would favor formation of rod-like fibrils as presence of the two viral protein fragments leads to an increased exposure of residues E46–A56, i.e., of the segment that forms the binding interface of the protofibrils in the rod fibril polymorph. Re-analyzing our old data we find for this segment in the simulations an average root-mean-square deviation (RMSD) to the corresponding chain segment in the rod fibril of about 5 Å for the control, but only 3 Å in presence of SK9, and 4 Å in presence of FI10. On the other hand, for the segment G68-A78 where in twister fibrils the protofibrils pack, the RMSD is with about 4 Å similar in control and presence of SK9 or FI10. Note that these averages have to be taken with a grain of salt as they are taken over all trajectories of Ref. 7 and 8, which differ in length and number of trajectories. However, even when taking these limitations into account, these averages show that rod-like or twister-like regions do appear in the ensemble of monomer conformations, and their frequency may depend on presence or absence of the viral protein fragments.

Our assumption is that dimers may form by binding of two αS chains at these two segments, leading to dimers that later would seed formation of either rod-like or twister-like αS. Assuming a threshold of 3.5 Å, we find that in the control simulation about 20% of conformations have rod-like segments, while in presence of SK9 or FI10 the frequency increases to about 50%. On the other hand, twister like segments are seen in about 50% of the conformation in control simulation or in presence of SK9, but only in about 40% of the conformations in simulations with FI10 present. These frequencies suggest that in presence of the viral protein fragments there is a higher probability for forming dimers by binding at the segment E46-A56 than seen in the control, but not for forming dimers that bind at the segment G68-A78. Albeit our monomer simulations suggest that presence of viral protein fragments increases the probability for aggregates binding at the same segment as in rod-like fibrils, we ignored this propensity difference in the monomer distributions (already reported by us in earlier work^8^), when setting up our dimer simulations. In order to avoid any biasing we generated instead model dimer configurations for both binding patterns. Using the procedure described in the method part, we selected the two conformations from our previous simulations (control and such generated in presence of FI10 or SK9) that had the lowest RMSD to either the Rod or the Twister segment. This procedure let to two model dimers bound at either the Rod binding site (E46-A56) or at the Twister binding site (G68-A78). For a third dimer we choose as chains the monomers with the highest strand content found in the earlier simulations. Sketches of the dimer models with and without viral protein fragments are shown in **Figure 1**. Two trajectories were followed over 500ns for each of the three dimer systems, both in presence of FI10 or SK9 and as control in absence of viral protein fragments. Technical details on all simulated systems are listed in **Table 1** and atomic coordinates of initial and final configurations are provided for each of the 18 trajectories as downloadable **supplementary information**.

**Figure 1:**
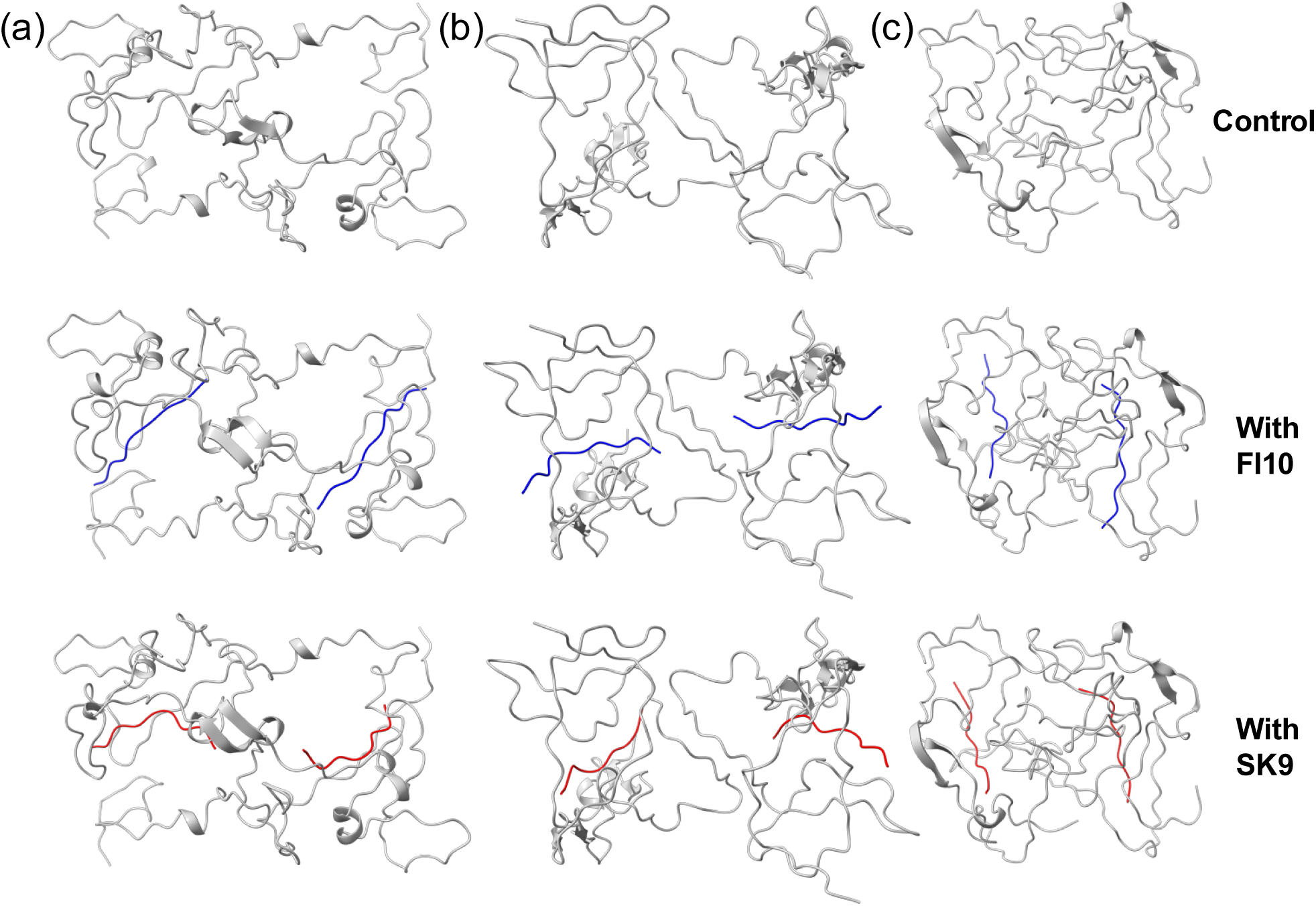
Sketches of the start configurations of our dimer simulations, with the Rod Binding dimers shown in (a), Twister Binding dimer in (b) and Beta Strand dimers in (c). The upper row shows the control, in the middle row is FI10 (in blue) binding to the dimers, and in the bottom SK9 (in red) binding to the dimers.

**Table I:**
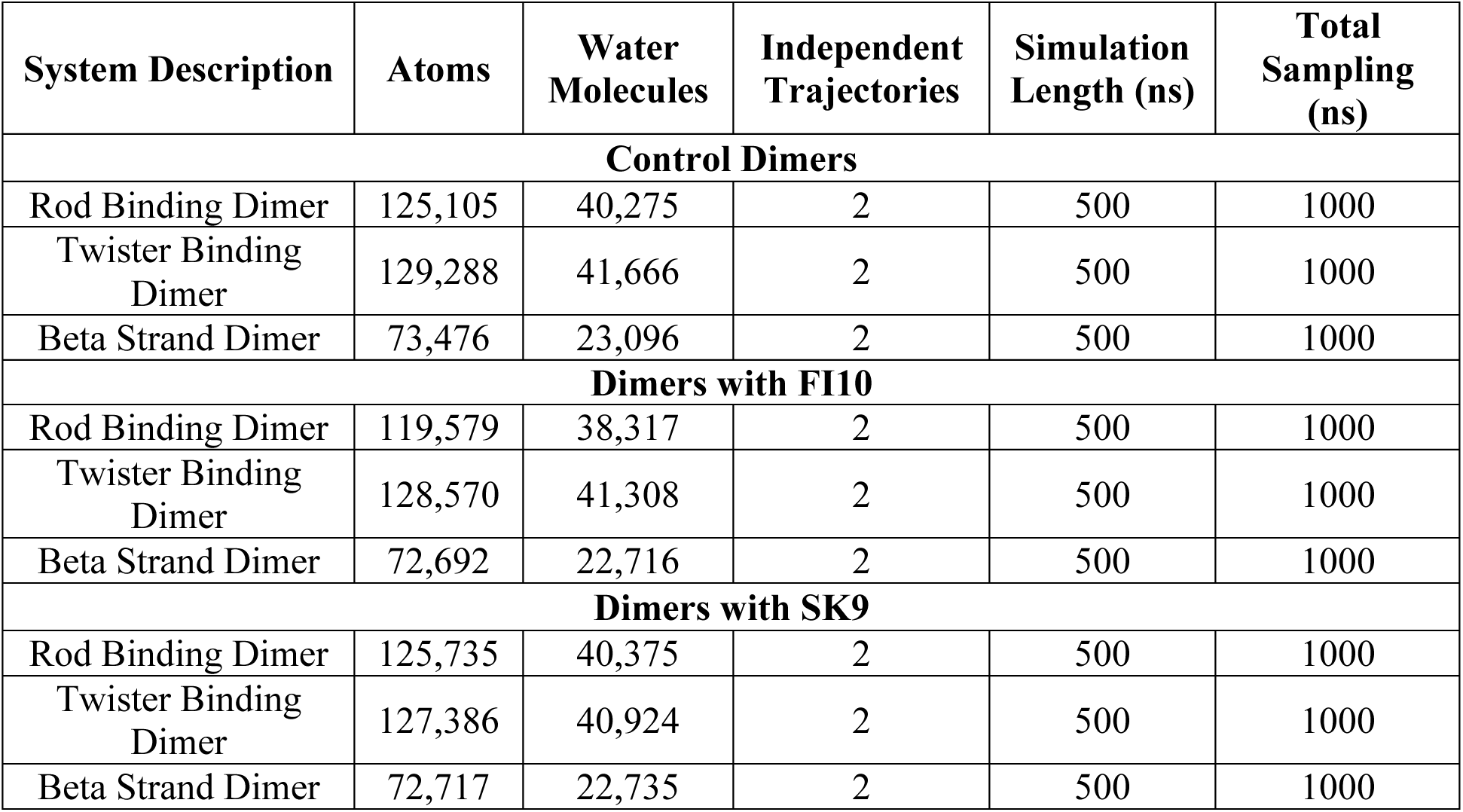
Simulated systems.

| System Description | Atoms | Water Molecules | Independent Trajectories | Simulation Length (ns) | Total Sampling (ns) |
| --- | --- | --- | --- | --- | --- |
| **Control Dimers** |  |  |  |  |  |
| Rod Binding Dimer | 125,105 | 40,275 | 2 | 500 | 1000 |
| Twister Binding Dimer | 129,288 | 41,666 | 2 | 500 | 1000 |
| Beta Strand Dimer | 73,476 | 23,096 | 2 | 500 | 1000 |
| **Dimers with FI10** |  |  |  |  |  |
| Rod Binding Dimer | 119,579 | 38,317 | 2 | 500 | 1000 |
| Twister Binding Dimer | 128,570 | 41,308 | 2 | 500 | 1000 |
| Beta Strand Dimer | 72,692 | 22,716 | 2 | 500 | 1000 |
| **Dimers with SK9** |  |  |  |  |  |
| Rod Binding Dimer | 125,735 | 40,375 | 2 | 500 | 1000 |
| Twister Binding Dimer | 127,386 | 40,924 | 2 | 500 | 1000 |
| Beta Strand Dimer | 72,717 | 22,735 | 2 | 500 | 1000 |

We start our analysis of these trajectories by looking into the stability of the binding of FI10 and SK9 to the three dimer models. For this purpose, we show in **Figure 2a** the total number of contacts between the two FI10 segments and the respective αS dimer models as a function of time, and in **Figure 2b** the corresponding plots for SK9. For FI10, we observe for all three dimer conformations a decrease in the number of contacts, indicating a de-stabilization, with the loss most pronounced for strand-dimers. We remark that binding of both FI10 and SK9 is not concentrated to certain pockets but rather unspecific to the residues of the αS dimers, and contacts between the viral protein fragments and the dimers may form and decay. For instance, the number of contacts between SK9 and the twister dimer is similar between start and finish of the trajectories, but decreases in between by half before the SK9 later re-attaches. As the viral protein fragments can move over the dimer and even detach, we plot in addition in **Figure 2c** and **2d** also the number of native contacts as function of time, that is the number of contacts that exist already at start. Here, we see in all cases a rapid decay, but with more native contacts surviving for SK9 binding to the dimers than for FI10 binding to the dimers. Initially, all models showed approximately 95 contacts between FI10 and the dimers. By the end of the simulation, only 5, 2, and 4 native contacts remained for the Rod Binding, Twister Binding, and Beta Strand dimers, respectively. In contrast, dimers with SK9 started with around 78 contacts, retaining 5, 4, and 6 native contacts for the Rod Binding, Twister Binding, and Beta Strand dimers at the end. The log-log plot in the insets shows that the loss of native contacts can be described by a power-law, indicative of diffusive motion of the viral fragments over the dimer surface.

**Figure 2:**
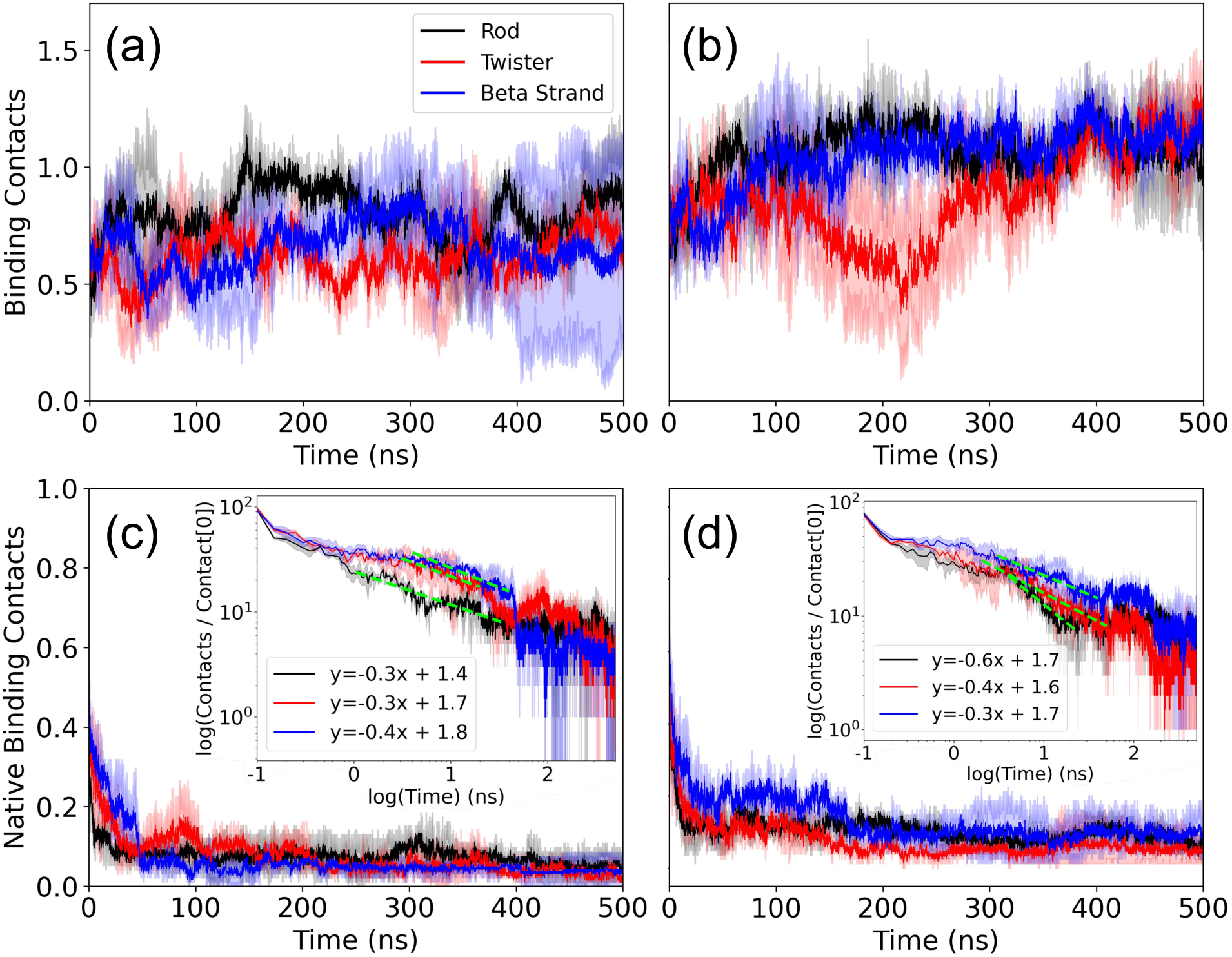
Time series of the number of contacts between the dimers and (a) the FI10 and (b) the SK9 viral protein fragment. Shown are averages over two trajectories, and the numbers normalized such that they are unity at start (t=0). In (c) and (d) we show the corresponding plots for the number of native contacts (that is contacts that are already observed at start), with the insets showing the time evolution of the same quantity in a log-log plot.

Combined, the four subfigures **2a** - **2d** indicate that SK9 binds stronger to all three dimer models than FI10, with the difference largest for the Beta Strand dimer and smallest for the Rod Binding dimer. Note that the curves in the four sub-plots have reached a plateau in the last 100 ns, and for this time interval, we find that SK9 residues form about 5 contacts with αS chains in any of the three dimer models while the numbers are with about 3 (Twister Binding) and 4 (Rod Binding and Beta Strand) smaller for FI10. Probabilities for SK9 or FI10 residues to form at least one contact with αS chains, evaluated over the last 100 ns and both trajectories, are given in the **Table 2** and show again a broad distribution, with stronger binding of residues of SK9 to the dimers than seen for FI10, and binding of FI10 residues especially weak to the Beta Strand dimer. When using the method of Ref. 21we find as binding free energies for FI10 −96 kJ/mol (Rod Binding), −86 kJ/mol (Twister Binding) and −29 kJ/mol (Beta Strand). For SK9 the corresponding values are −127 kJ/mol (Rod Binding), −111 kJ/mol (Twister Binding) and −72 kJ/mol (Beta Strand). Hence, our data suggest that SK9 will affect the αS dimers more than FI10, and that the effect of SK9 is strongest for the Rod Binding dimer.

**Table 2:** Probabilities for SK9 or FI10 residues to form at least one contact with αS chains, evaluated over the last 100ns. Values are averaged over each fragment and both trajectories.

|  | FI10 |  |  | SK9 |  |  |
| --- | --- | --- | --- | --- | --- | --- |
| Residue | Rod Binding Dimer | Twister Binding Dimer | Beta Strand Dimer | Rod Binding Dimer | Twister Binding Dimer | Beta Strand Dimer |
| 1 | 0.86 | 0.85 | 0.75 | 0.88 | 0.95 | 0.99 |
| 2 | 0.80 | 0.88 | 0.72 | 0.99 | 0.99 | 0.95 |
| 3 | 0.89 | 0.82 | 0.70 | 0.99 | 0.99 | 1 |
| 4 | 0.88 | 0.77 | 0.67 | 0.99 | 0.99 | 0.99 |
| 5 | 0.99 | 0.88 | 0.50 | 0.99 | 0.99 | 1 |
| 6 | 0.99 | 0.93 | 0.47 | 0.90 | 0.91 | 0.94 |
| 7 | 0.98 | 0.99 | 0.72 | 0.97 | 0.94 | 0.99 |
| 8 | 0.90 | 0.98 | 0.71 | 0.99 | 0.88 | 0.85 |
| 9 | 0.88 | 0.88 | 0.78 | 0.98 | 0.91 | 0.87 |
| 10 | 0.67 | 0.83 | 0.75 | - | - | - |

How is the binding of FI10 or SK9 altering the evolution of the dimers? The first quantity that we considered is the radius of gyration (R_g_), a measure of the dimer extension, and the average sheetness, i.e., the percentage of residues that were identified as strand-like. Averages of these and other quantities, taken over the final 100 ns and both trajectories are shown for system in **Table 3** for Rod Binding, Twister Binding, and for Beta Strand dimers. At start (t=0) we find a R_g_ of about 2.8 nm and a sheetness of 6% for the Rod Binding dimer, 2.6 nm and 14% for Twister Binding dimers, and 2.2 nm and 20% for Beta Strand dimers. Averaged over the last 100 ns the R_g_ values decrease for Twister Binding dimers by 0.2 nm in the control and 0.1 nm in presence of FI10 and 0.5 nm in the presence of SK9, while the sheetness stays unchanged in all three cases. Similarly, the R_g_ values decrease for Rod Binding dimers by about 0.1 nm in the control and increase by 0.1 nm or stay unchanged in the presence of FI10 or SK9, however, the sheetness increases by 5% in the control while only by 1% in presence of the viral protein fragments. The situation is different for the Beta Strand dimers, where the R_g_ increases by 0.1 nm for the control and 0.6 nm in presence of FI10, but only stays unchanged in presence of SK9. At the same time, the sheetness decreases here by 7% in all three cases. Hence, presence of the viral fragments has little or no effect on the compactness or the sheetness of the dimer conformations, with only some minor stabilization seen for dimers with high strandness of the chains.

**Table 3:**
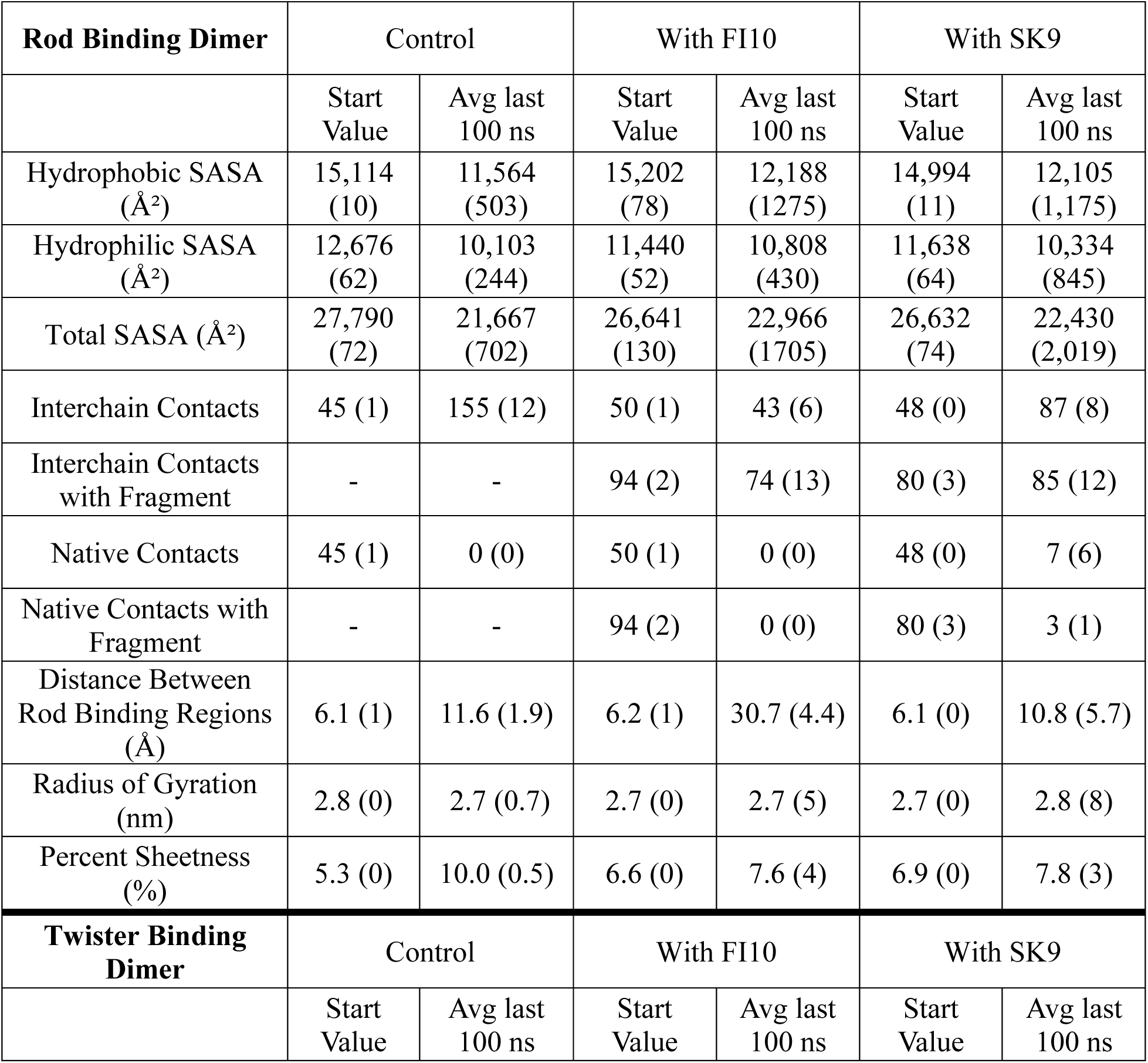
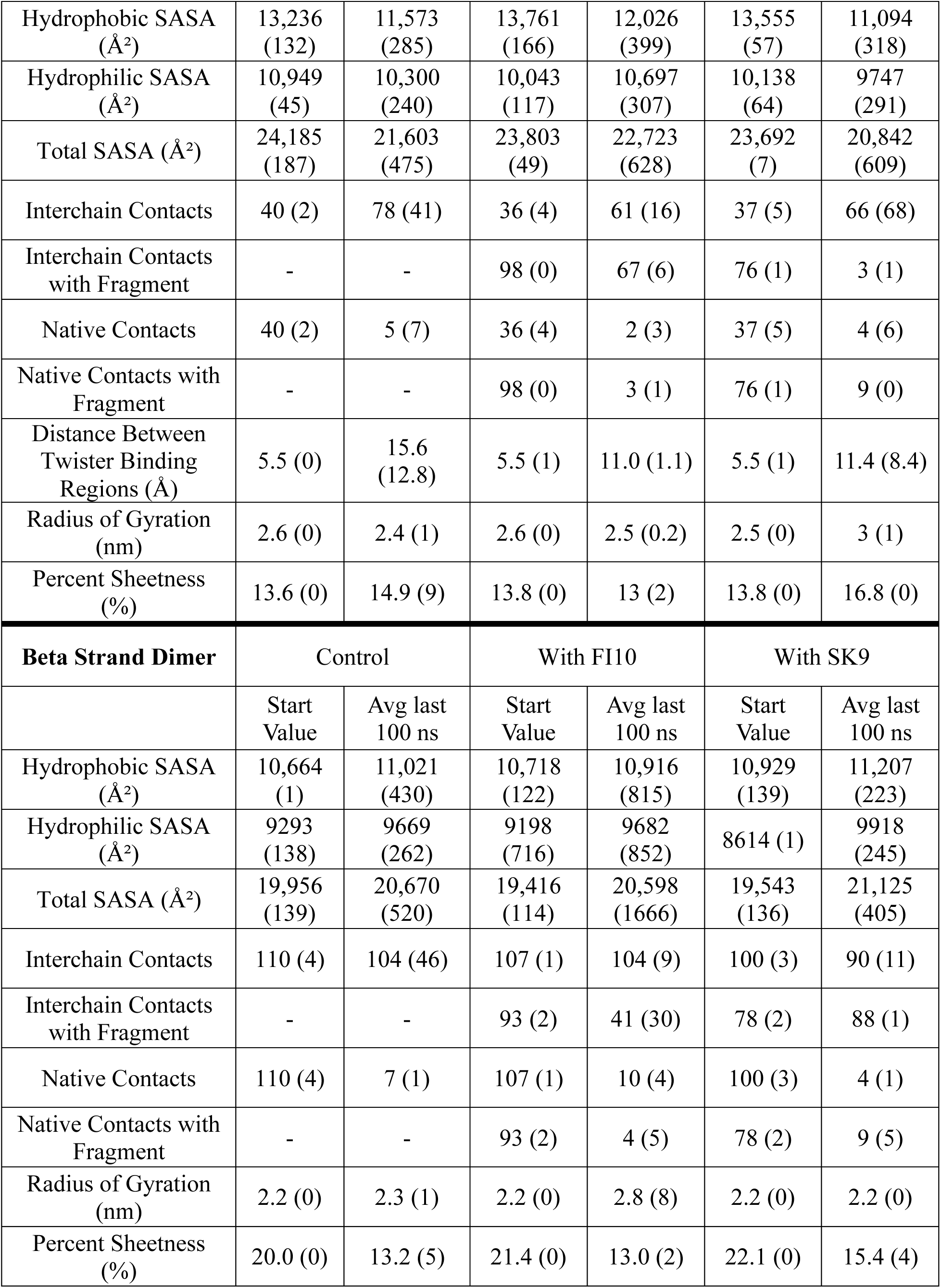
Various quantities measured over the last 100 ns in simulations of Rod Binding, Twister Binding, and Beta Strand dimers in presence of either FI10 or SK9, or in absence of the viral protein fragments. Averages over two trajectories for each system are shown, with standard deviations in parenthesis.

| **Rod Binding Dimer** | Control |  | With FI10 |  | With SK9 |  |
| --- | --- | --- | --- | --- | --- | --- |
|  | Start Value | Avg last 100 ns | Start Value | Avg last 100 ns | Start Value | Avg last 100 ns |
| Hydrophobic SASA ($\text{\AA}^2$) | 15,114 (10) | 11,564 (503) | 15,202 (78) | 12,188 (1275) | 14,994 (11) | 12,105 (1,175) |
| Hydrophilic SASA ($\text{\AA}^2$) | 12,676 (62) | 10,103 (244) | 11,440 (52) | 10,808 (430) | 11,638 (64) | 10,334 (845) |
| Total SASA ($\text{\AA}^2$) | 27,790 (72) | 21,667 (702) | 26,641 (130) | 22,966 (1705) | 26,632 (74) | 22,430 (2,019) |
| Interchain Contacts | 45 (1) | 155 (12) | 50 (1) | 43 (6) | 48 (0) | 87 (8) |
| Interchain Contacts with Fragment | - | - | 94 (2) | 74 (13) | 80 (3) | 85 (12) |
| Native Contacts | 45 (1) | 0 (0) | 50 (1) | 0 (0) | 48 (0) | 7 (6) |
| Native Contacts with Fragment | - | - | 94 (2) | 0 (0) | 80 (3) | 3 (1) |
| Distance Between Rod Binding Regions ($\text{\AA}$) | 6.1 (1) | 11.6 (1.9) | 6.2 (1) | 30.7 (4.4) | 6.1 (0) | 10.8 (5.7) |
| Radius of Gyration (nm) | 2.8 (0) | 2.7 (0.7) | 2.7 (0) | 2.7 (5) | 2.7 (0) | 2.8 (8) |
| Percent Sheetness (%) | 5.3 (0) | 10.0 (0.5) | 6.6 (0) | 7.6 (4) | 6.9 (0) | 7.8 (3) |
| **Twister Binding Dimer** | Control |  | With FI10 |  | With SK9 |  |
|  | Start Value | Avg last 100 ns | Start Value | Avg last 100 ns | Start Value | Avg last 100 ns |
| Hydrophobic SASA (Å<sup>2</sup>) | 13,236 (132) | 11,573 (285) | 13,761 (166) | 12,026 (399) | 13,555 (57) | 11,094 (318) |
| Hydrophilic SASA (Å<sup>2</sup>) | 10,949 (45) | 10,300 (240) | 10,043 (117) | 10,697 (307) | 10,138 (64) | 9747 (291) |
| Total SASA (Å<sup>2</sup>) | 24,185 (187) | 21,603 (475) | 23,803 (49) | 22,723 (628) | 23,692 (7) | 20,842 (609) |
| Interchain Contacts | 40 (2) | 78 (41) | 36 (4) | 61 (16) | 37 (5) | 66 (68) |
| Interchain Contacts with Fragment | - | - | 98 (0) | 67 (6) | 76 (1) | 3 (1) |
| Native Contacts | 40 (2) | 5 (7) | 36 (4) | 2 (3) | 37 (5) | 4 (6) |
| Native Contacts with Fragment | - | - | 98 (0) | 3 (1) | 76 (1) | 9 (0) |
| Distance Between Twister Binding Regions (Å) | 5.5 (0) | 15.6 (12.8) | 5.5 (1) | 11.0 (1.1) | 5.5 (1) | 11.4 (8.4) |
| Radius of Gyration (nm) | 2.6 (0) | 2.4 (1) | 2.6 (0) | 2.5 (0.2) | 2.5 (0) | 3 (1) |
| Percent Sheetiness (%) | 13.6 (0) | 14.9 (9) | 13.8 (0) | 13 (2) | 13.8 (0) | 16.8 (0) |
| **Beta Strand Dimer** | Control |  | With FI10 |  | With SK9 |  |
|  | Start Value | Avg last 100 ns | Start Value | Avg last 100 ns | Start Value | Avg last 100 ns |
| Hydrophobic SASA (Å<sup>2</sup>) | 10,664 (1) | 11,021 (430) | 10,718 (122) | 10,916 (815) | 10,929 (139) | 11,207 (223) |
| Hydrophilic SASA (Å<sup>2</sup>) | 9293 (138) | 9669 (262) | 9198 (716) | 9682 (852) | 8614 (1) | 9918 (245) |
| Total SASA (Å<sup>2</sup>) | 19,956 (139) | 20,670 (520) | 19,416 (114) | 20,598 (1666) | 19,543 (136) | 21,125 (405) |
| Interchain Contacts | 110 (4) | 104 (46) | 107 (1) | 104 (9) | 100 (3) | 90 (11) |
| Interchain Contacts with Fragment | - | - | 93 (2) | 41 (30) | 78 (2) | 88 (1) |
| Native Contacts | 110 (4) | 7 (1) | 107 (1) | 10 (4) | 100 (3) | 4 (1) |
| Native Contacts with Fragment | - | - | 93 (2) | 4 (5) | 78 (2) | 9 (5) |
| Radius of Gyration (nm) | 2.2 (0) | 2.3 (1) | 2.2 (0) | 2.8 (8) | 2.2 (0) | 2.2 (0) |
| Percent Sheetiness (%) | 20.0 (0) | 13.2 (5) | 21.4 (0) | 13.0 (2) | 22.1 (0) | 15.4 (4) |

We find a clearer signal in our measurements of the solvent accessible surface area (SASA) in **Figure 3** which describes the hydration of the dimers. For the Rod Binding dimer (**Figure 3a -3c**) we find in all three cases a decreases of SASA of about 4000Å^2^ over the first 50 ns that continues for the control to a total loss of about 6000 Å^2^ at 500ns, while a plateau is reached in presence of SK9 (at a loss of about 4200 Å^2^) or FI10 (at a loss of about 3600 Å^2^). The difference to the control is especially visible for the SASA component resulting from hydrophilic residues. For these residues are 2500 Å^2^ lost in the control but only 600 Å^2^ for FI10 and 1300 Å^2^ for SK9. The situation is different for Twister Binding dimers (**Figure 3d-3f**) where the loss of SASA approaches a plateau after about 250ns for control (−3600 Å^2^) and SK9 (−2900 Å^2^), but increases again in presence of FI10, resulting in a final loss of only 1100 Å^2^. This effect is again mostly due to contributions from hydrophilic residues for which the SASA increases by about 650 Å^2^ in presence of FI10, but decreases by about 380 Å^2^ in presence of SK9 and 650 Å^2^ in the control. For Beta Strand dimers (**Figure 3g-3i**) is again a plateau reached only after about 250 ns, with little change in SASA for the control (+760 Å^2^), and a larger gain in presence of SK9 (+1600 Å^2^) or FI10 (+1200 Å^2^) resulting for SK9 mostly from hydrophilic residues (+1300 Å^2^). Hence, the viral protein fragments either minimize the loss of SASA for hydrophilic residues or even increase their exposure, with the effect stronger for FI10 than SK9 in Twister Binding and Rod Binding dimers.

**Figure 3:**
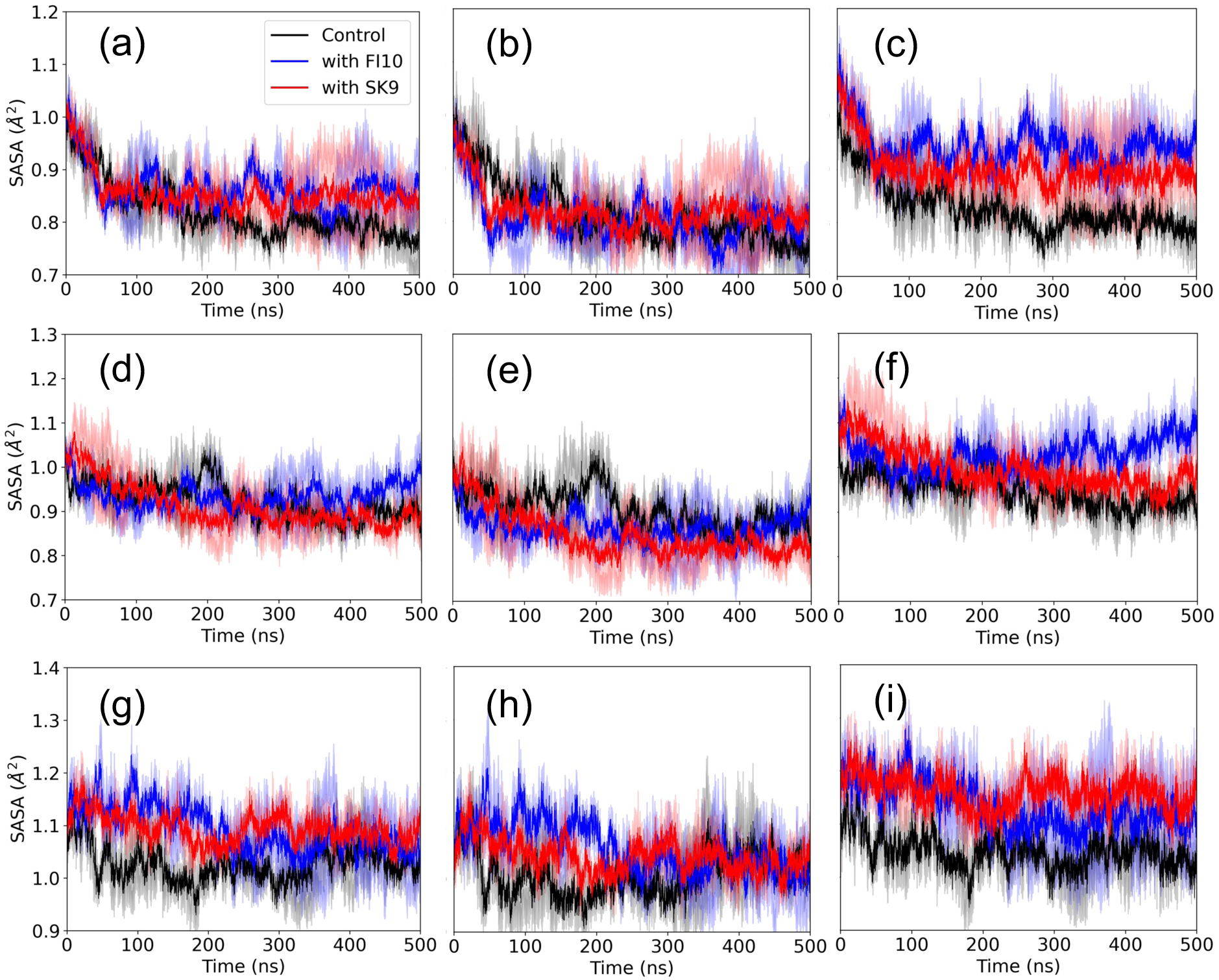
Solvent accessible surface area (SASA) as function of time for Rod Binding dimers (a-c), Twister Binding dimers (d-f) and Beta Strand dimers (g-i) for both control simulations and such in presence of either FI10 or SK9. Shown are averages over two independent trajectories, and the values divided by the one at start. Values calculated over all residues are shown in the left row, while the center row shows the values calculated only for hydrophobic residues and the right one the values calculated over hydrophilic residues.

The above discussed quantities describe global properties of the dimers. In order to understand in more detail how interaction with viral protein fragments affects the three dimer models we continue our analysis by looking into the number of interchain contacts between the two monomers in a dimer. The time evolution of this quantity, which is a measure for the stability of the dimer, is shown for the three systems in **Figure 4**. To allow for an easy comparison, we normalize again our data by dividing them through the respective start value. In all cases we observe a quick plateauing for the simulations with FI10 or SK9 interacting with dimers, while (with the exception of the Beta Strand dimers) this process can take more than 200 ns for the control. For all three dimer models we find that the number of such interchain contacts increases or stays similar to the start values (**Figure 4a** - **4c**) while at the same time the number of native interchain contacts (that is such contacts seen already at start, shown in **Figure 4d** - **4f**) decreases rapidly. The increase in the total number of interchain contacts is smaller in presence of SK9 or FI10 than in the control, see **Table 3**, with the effect stronger for FI10 than for SK9 and best seen for the Rod Binding dimer. On the other hand, the decrease of native contacts is in all three dimer models similar in control and FI10, but (with the exception of the beta-strand dimer) lower for SK9. For instance, in the Rod Binding dimer no native contacts are found in the final 100 ns of control simulations or such in presence of FI10, but about 10 in such where SK9 is present. Hence, presence of SK9 or FI10 stabilizes the original Rod Binding or Twister Binding dimer conformations, in this way delaying the decay of the respective start conformations that allows the two chains to move relative to each other and to form additional contacts. The protecting effect seems to be stronger for SK9 in the Rod Binding dimer while for the Twister Binding dimer both viral protein fragments have a similar effect and no such stabilization is seen in Beta Strand dimers.

**Figure 4:**
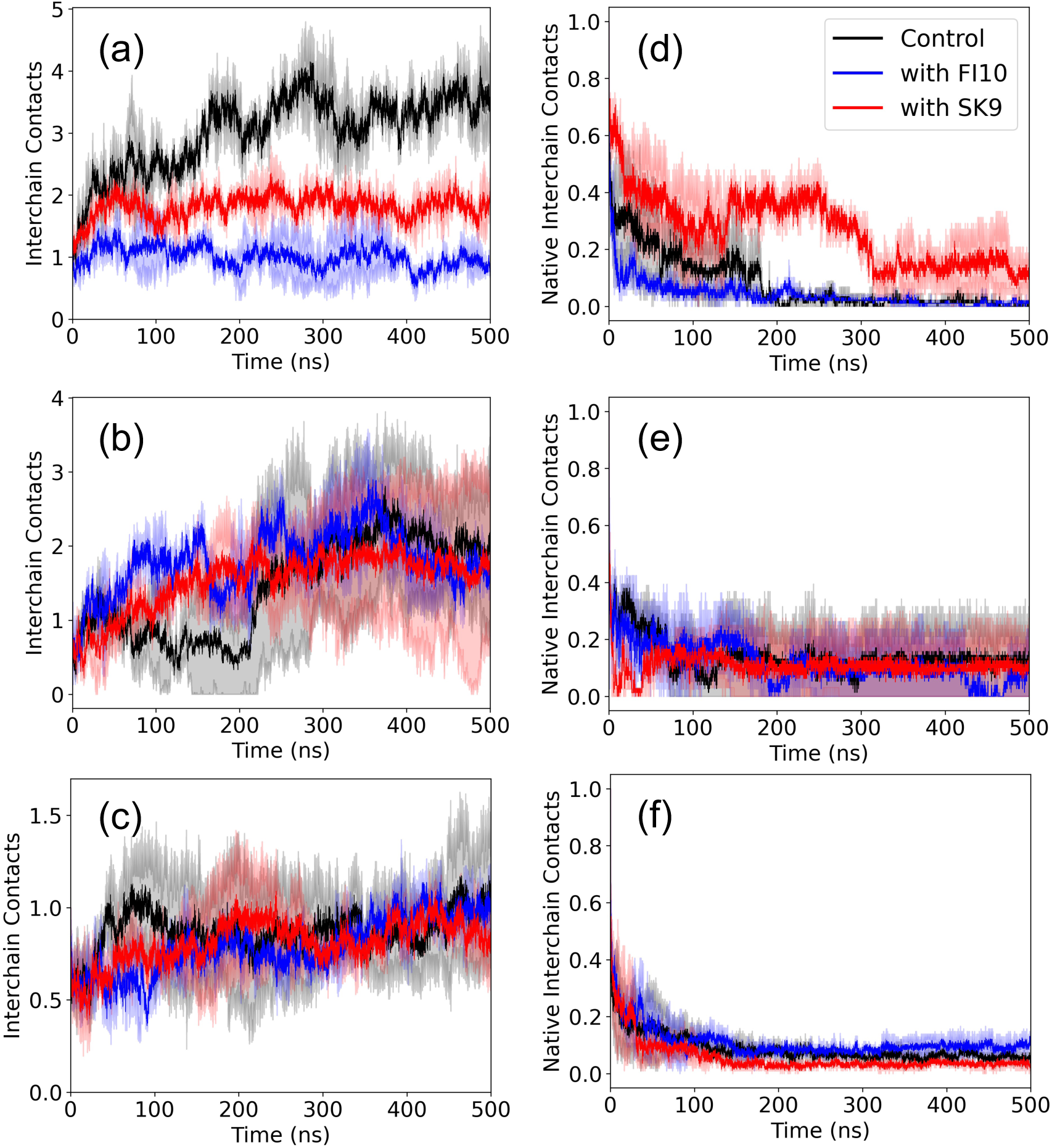
Number of interchain contacts (normalized to a start value of unity) between αS chains in the Rod Binding (upper row), Twister Binding (middle row), and Beta Strand (lower row) dimers, both in presence or absence of FI10 or SK9. Shown are averages over two independent trajectories. The plots in the right column shows the corresponding numbers when only contacts are counted that exist already at start.

The more pronounced effect of SK9 and FI10 on the Rod Binding and Twister Binding dimers when compared to the Beta Strand dimers may be because the Rod Binding and Twister Binding dimers have defined interaction interfaces. In order to test this hypothesis, we also look specifically into the interchain contacts at these interfaces (the segment of residues E46-A56 for the Rod Binding dimers and G68-A78 for Twister Binding dimers). We show in **Figure 5a** and **5b** again the number of total contacts and the number of native contacts for Rod Binding dimers, and in **Figure 5d** and **5e** for Twister Binding dimers, normalized to a start value of one for more easy comparison. Average values taken over the last 100 ns are listed in **Table 4**.

**Figure 5:**
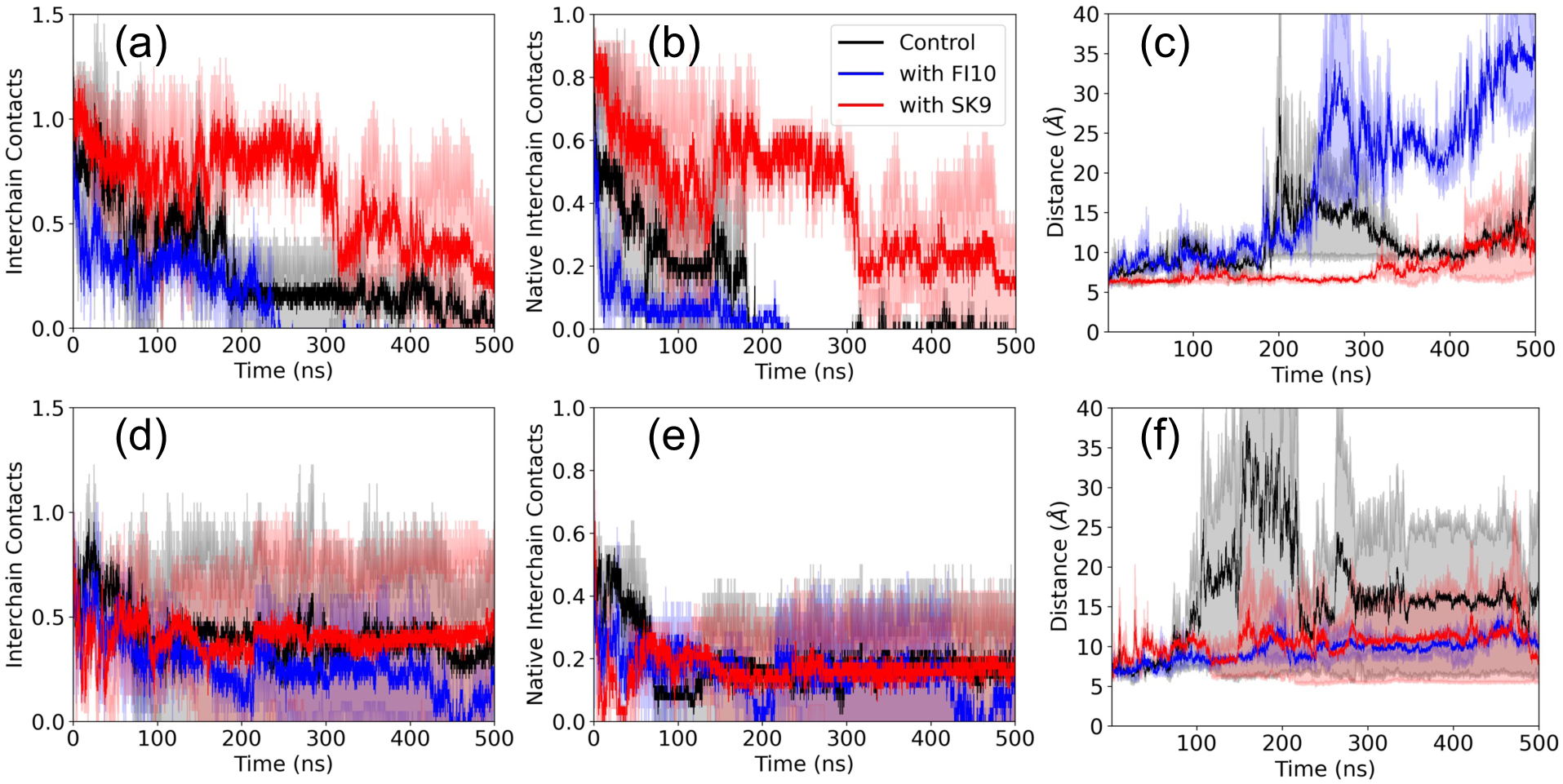
Number of interchain contacts (normalized to a start value of unity and averaged over two independent trajectories) between αS chains in the respective binding regions of Rod Binding dimers (upper row) and Twister Binding dimers (lower row) as function of time for the control and in presence of either FI10 or SK9. The right column shows the total number of such contacts, while the center column shows the number of native contacts, i.e., such already present at start. The right column shows the distance between the two chains at the respective binding segment.

**Table 4:**
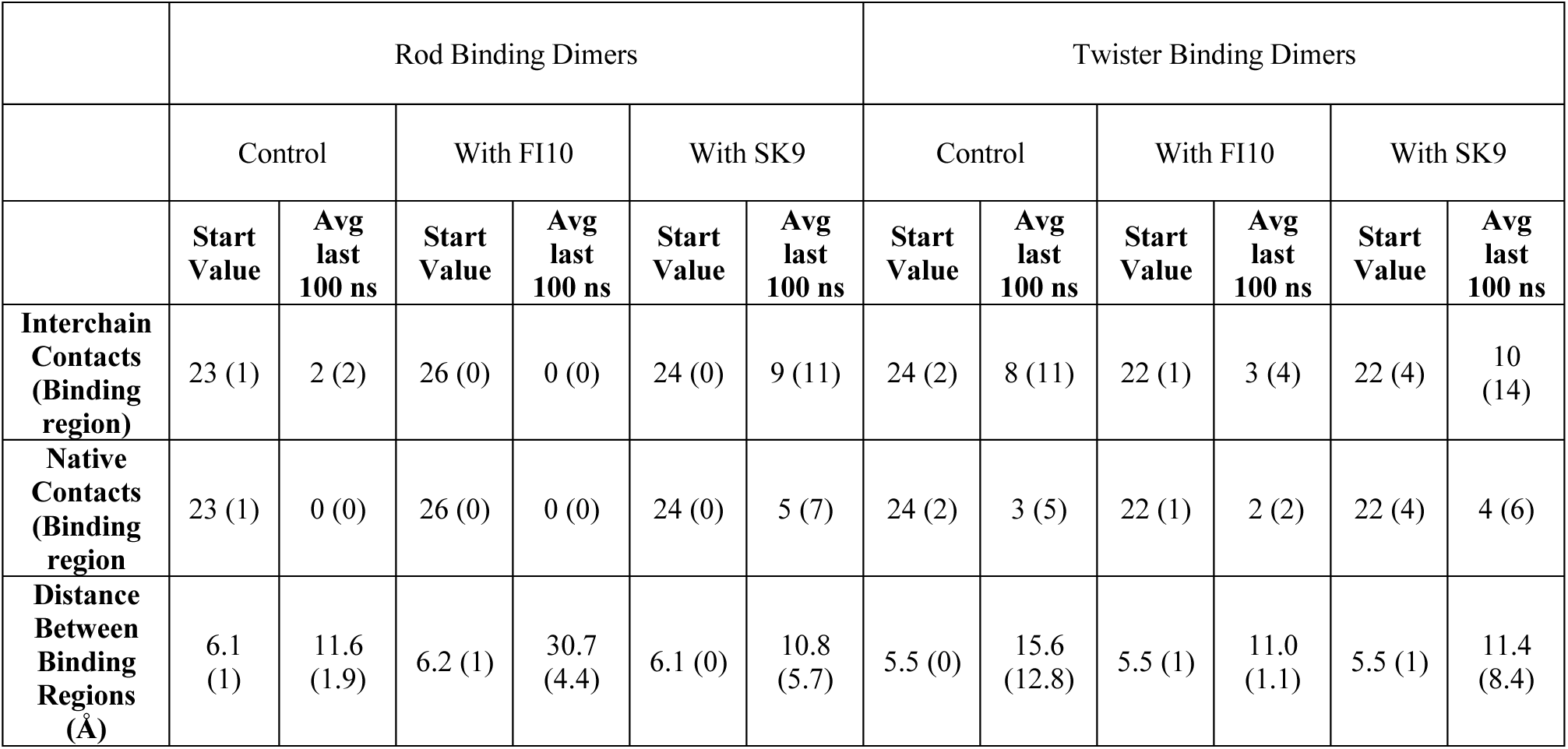
Interchain contacts between the binding regions of Rod Binding or Twister Binding dimers in presence or absence of the viral protein fragments SK9 and FI10.

|  | Rod Binding Dimers |  |  |  |  |  | Twister Binding Dimers |  |  |  |  |  |
| --- | --- | --- | --- | --- | --- | --- | --- | --- | --- | --- | --- | --- |
|  | Control |  | With FI10 |  | With SK9 |  | Control |  | With FI10 |  | With SK9 |  |
|  | Start Value | Avg last 100 ns | Start Value | Avg last 100 ns | Start Value | Avg last 100 ns | Start Value | Avg last 100 ns | Start Value | Avg last 100 ns | Start Value | Avg last 100 ns |
| **Interchain Contacts (Binding region)** | 23 (1) | 2 (2) | 26 (0) | 0 (0) | 24 (0) | 9 (11) | 24 (2) | 8 (11) | 22 (1) | 3 (4) | 22 (4) | 10 (14) |
| **Native Contacts (Binding region)** | 23 (1) | 0 (0) | 26 (0) | 0 (0) | 24 (0) | 5 (7) | 24 (2) | 3 (5) | 22 (1) | 2 (2) | 22 (4) | 4 (6) |
| **Distance Between Binding Regions (Å)** | 6.1 (1) | 11.6 (1.9) | 6.2 (1) | 30.7 (4.4) | 6.1 (0) | 10.8 (5.7) | 5.5 (0) | 15.6 (12.8) | 5.5 (1) | 11.0 (1.1) | 5.5 (1) | 11.4 (8.4) |

While the number of interchain contacts calculated over the full lengths of the chains increases in the two dimer models along the trajectories, this is not the case for the number of contacts between the two chains at the defined rod or twister binding interfaces. In both dimer models do these numbers decrease rapidly, and the differences between the number of such contacts, and that of the native contacts, measured over the final 100 ns is small, see **Table 2** and **Table 4**. This indicates that dissolution of native contacts is often not compensated by formation of new contacts, indicating a separation of the chains at the respective interface.

This is also seen in **Figure 5c** and **5f**, where the distance between the two interfaces grows with time in all systems. For the Rod Binding dimers we find that SK9 but not FI10 stabilizes binding at this interface. While in the control and in presence of FI10 the distance between the two interfaces increase from the start and doubles its values at about 180 ns for the control, and at about 250 ns in presence of FI10, it does not grow in presence of SK9 for about 320 ns, and even later the distance stays always lower than for the control and even more for FI10 where the rod dimer appears to dissociate. Note that the time points where the distance between the rod interfaces starts increasing rapidly also corresponds with the times where rapid loss of interchain contacts is seen. On the other hand, in the Twister Binding dimer the number of interchain contacts connecting the corresponding regions differs little between control and trajectories where SK9 or FI10 are present, but the relative loss of contacts is smaller than for the Rod Binding dimer. However, the separation between the interfaces grows similar as for the Rod Binding dimer, with the growth smaller in presence of FI10 and SK9 than in the control. For instance, this distance reaches for the control at around 80 ns a value that is double that of its start values, and is settling over the last 100 ns at around 15 Å; while in presence of SK9 and FI10 it takes about 180 ns before the initial distance has doubled, and afterwards stays almost constant until reaching final values of about 10 Å. As for the Rod Binding dimer is the increase of distance between the binding interfaces connected with the loss in native interchain contacts.

## III. CONCLUSIONS

Using long molecular dynamics simulations, we have studied the effect of two short SARS-COV-2 fragments, SK9 and FI10, on the stability of three αS-dimer models that we regard as potential seeds for the neurotoxic aggregates associated with the pathogenesis of Parkinson disease. We find that SK9 binds to all three models stronger than FI10 does, and moves less over the surface of the dimers, even increasing the number of contacts to the dimers, especially the Beta Strand dimer. Both viral protein fragments have little effect on the time evolution of the number of interchain contacts in the Beta Strand dimer but compared to the control reduce the loosening and loss of secondary structure of the conformations, and increase its solvent accessible surface, both for hydrophobic and hydrophilic residues. Hence, both fragments stabilize the Beta Strand dimer, but the effect is stronger for SK9 which also binds more strongly to this dimer model.

For Rod Binding and Twister Binding dimers we do not see any effect of the viral fragments on the secondary structure contacts, and even a larger loosening of the dimers that goes together with a smaller increase in the number of interchain contacts than seen in the control. On the other hand, the decrease of native contacts is slightly less for FI10 and SK9 than in the control, with the effect more pronounced for SK9. In the Rod Binding dimer are no native contacts found in the final 100 ns of control simulations or such in presence of FI10, but about 10 contacts when SK9 is present. This suggests that SK9, and to a smaller degree also FI10, stabilize the original conformation of the Rod Binding and Twister Binding dimers, delaying movement of the two chains relative to each other that could lead to forming additional contacts. The protecting effect especially of SK9 seems to be stronger for the Rod Binding dimer, where it leads to a reduced decrease in solvent accessible surface area that again results from both hydrophobic and hydrophilic residues. Note that for the Twister Binding dimer the decrease in SASA seen in the control is reversed for FI10 but not for SK9. This is because of a strong increase in SASA for hydrophilic residues. An increase (or smaller decrease) of exposure of hydrophilic residues in presence of FI10 (and to a lesser extend also of SK9) is observed in both dimer models and seems to be one of the mechanisms for the effect of the protein fragments on the dimers. Rod Binding and Twister Binding dimers were designed to have binding interfaces at the segments observed in two experimentally resolved fibril polymorphs. Measuring the number of interchain contacts and distance between the respective segments, we find that the presence of SK9 or FI10 stabilizes both twister and rod interface, but the effect is more pronounced for the Rod Binding dimer. Here we see a strong protective effect by SK9 but none by FI10 whereas in the control the chains even separate.

In summary, our results indicate that the two viral protein fragments may stabilize αS-dimer, potentially allowing them to seed neurotoxic aggregates. The extend and mechanism depends both on the fragment and the dimer model, but our results suggest that SK9 has a larger effect than FI10, stabilizing existing structural elements (such as binding interfaces or secondary structure), and encouraging exposure of hydrophilic residues.

## IV. MATERIALS AND METHODS

### A. System Preparation

Our study aims to understand αS aggregation by investigating αS dimers as being the simplest oligomers. We use all-atom molecular dynamics (MD) simulations to study the change in conformational ensemble of αS dimers induced by the presence of the ten-residue segment _194_FKNIDGYFKI_203_ (FI10) of the spike protein and the nine-residue segment _54_SFYVYSRVK_62_ (SK9) of the envelope protein from SARS-COV-2. The initial configurations for dimer models were obtained by docking homogenous monomers taken from previous MD simulations of the αS monomer.^7, 8^ The equilibrated monomer conformations were selected for the Rod Binding and Twister Binding dimers based on structural similarity of the protofilament interfaces to the same regions in cryo-EM structures of the corresponding Rod and Twister αS fibrils, as measured by root-mean-square deviation (RMSD). Specifically, the selection was based on the protofibril binding region (E46-A56) in the Rod polymorph (PDB ID: 6CU7) and (G68-A78) in the Twister polymorph (PDB ID: 6CU8).^9^ For the Beta Strand dimer, monomers were chosen from conformations with the highest percentage of sheetness as determined by VMD and the STRIDE algorithm.^10^ Dimers were generated by docking identical monomers by HADDOCK program using standard protein-protein parameters.^11, 12^ For the Rod and Twister Binding dimers the protofibril binding regions were selectively docked to form dimers with these regions in contact. In all models, we capped the N- and C-terminal of each αS monomer in the dimer models with a NH_3_^+^ and a COO^−^ group.

Simulations starting from the αS dimer models described above serve as control for simulations where in addition also FI10 or SK9 peptides are present. Initial configurations of FI10 and SK9 were prepared as described in earlier work,^7, 8^ with the N- and C-terminal FI10 capped by a NH_3_^+^ and -COO^−^ group, respectively, and, in order to stay consistent with earlier work, SK9 capped by a NH_3_^+^ and -CONH_2_ group. To produce the initial conformations for our simulations, we docked two FI10 and SK9 segments to each dimer at binding sites predicted by HADDOCK when using standard protein-peptide parameters.^11, 12^ A single fragment was docked to each monomer in the dimer producing symmetrical initial configurations. Note that the viral protein segments are not fixed but can move freely throughout the simulations and may detach from the αS chains. The so obtained start configurations are shown in **Figure 1**, and their atomic coordinates are provided as downloadable **supplementary information**.

### B. General Simulation Protocol

The dimer systems were simulated with the with the GROMACS package 2022 package^13^ using the CHARMM 36m all-atom force-field^14^ with TIP3P explicit water.^15^ Hydrogen atoms were added with the *pdb2gmx* module of the GROMACS suite.^13^ The start configurations for each system were placed in the center of a cubic box with periodic boundary conditions and an edge-length in each direction of 10.56 nm for the Rod Binding dimer, 10.81 nm for the Twister Binding dimer, and 10.74 nm for the Beta Strand dimer. The simulation boxes were solvated with water molecules, and Na^+^ and Cl^−^ ions were added at a physiological ion concentration of 150 mM NaCl to neutralize the system. **Table I** lists the total number of atoms and number of water molecules in each system. Energy minimization occurred by steepest decent for up to 50,000 steps, followed by short molecular dynamics simulation at 310K for 200 ps at constant volume and an additional 200ps at constant pressure (1 atm), constraining the positions of heavy atoms with a force constant of 1000 kJ mol^−1^ nm^−2^.

During the trajectories the temperature was held constant at 310 K by a v-rescale thermostat^16^ with a coupling constant of 0.1 ps and pressure was maintained at a constant 1 atm by using the Parrinello-Rahman barostat^17^ with a pressure relaxation time of 2 ps. The SETTLE algorithm^18^ keeps water molecules rigid, and protein bonds involving hydrogen atoms are restrained to their equilibrium length with the LINCS algorithm.^19^ We used a time step of 2 fs for integrating the equations of motion. Because of periodic boundary conditions, we used the particle-mesh Ewald (PME) technique to calculate the long-range electrostatic interactions, with a real-space cutoff of 12 Å and a Fourier grid spacing of 1.6 Å. Short-range van der Waal interactions were truncated at 12 Å, with smoothing starting at 10.5 Å. In this study, we considered two trajectories of 500 ns for each model with different initial velocity distributions.

### C. Trajectory Analysis

GROMACS tools^13^ and VMD are used to analyze trajectories. VMD is used to visualize conformations.^20^ GROMACS tools are used to calculate root-mean-square deviation (RMSD), root-mean-square-fluctuation (RMSF), radius of gyration (RGY), solvent accessible surface area (SASA), distance between binding regions, and contact frequencies. The calculation of SASA relied on a spherical probe of 1.4 Å radius. Contacts are defined by a cutoff 4.5 Å in the closest distance between heavy atoms in a residue pair. Residue-wise secondary structure propensity is calculated using VMD and the STRIDE algorithm.^10^ Helicity and sheetness are defined as the percentage of residues with alpha-helical (i.e., assigned a “H” by STRIDE) or beta-strand (an “E” for extended configuration or a “B” for an isolated bridge) secondary structure in every frame of the trajectory. Binding free energies are calculated with the method described in Ref. 21 where one has to account for the difference in chain lengths when calculating the concentration *C_sim_*.

## Supporting information

Folder with atomic coordinates of start and final configurations of all trajectories in PDB format

## SUPPORTING INFORMATION

The Supporting Information is available free of charge at URL

- Start and final configurations of all trajectories in PDB format as text files in a compressed folder *aS_Configurations-Supporting_Information.zip*

## AUTHOR INFORMATION

### Author Contributions

**Lucy M Coleman:** Formal analysis (equal); Investigation (equal); Visualization (lead); Writing – original draft (equal); **Ulrich H.E. Hansmann:** Conceptualization (lead); Funding acquisition (lead); Resources (lead); Supervision (lead); Writing – original draft (equal); Writing – review and editing (lead).

## ACKNOWLEDGMENT

The simulations in this work were done using the SCHOONER cluster of the University of Oklahoma, XSEDE resources allocated under grant MCB160005 (National Science Foundation). We thank Andrew D Chesney for help at an early stage of this study with setting up and running simulations.

## DATA AVAILABILITY

The data that support the findings of this study are available in the Supporting Information and are publicly accessible at https://github.com/ouhansmannlab/alpha_synuclein.git

## REFERENCES

1 Spillantini, M. G.; Schmidt, M. L.; Lee, V. M.-Y.; Trojanowski, J. Q.; Jakes, R.; Goedert, M. α-synuclein in Lewy bodies. Nature 1997, 388 (6645), 839–840. DOI: 10.1038/42166.

2 Meade, R. M.; Fairlie, D. P.; Mason, J. M. Alpha-synuclein structure and Parkinson’s disease - lessons and emerging principles. Mol. Neurodegener. 2019, 14 (1), 29. DOI: 10.1186/s13024-019-0329-1.

3 Leta, V.; Boura, I.; van Wamelen, D. J.; Rodriguez-Violante, M.; Antonini, A.; Chaudhuri, K. R. Covid-19 and Parkinson’s disease: Acute clinical implications, long-COVID and post-COVID-19 parkinsonism. Int Rev Neurobiol 2022, 165, 63–89. DOI: 10.1016/bs.irn.2022.04.004.

4 Semerdzhiev, S. A.; Fakhree, M. A. A.; Segers-Nolten, I.; Blum, C.; Claessens, M. Interactions between SARS-CoV-2 N-Protein and alpha-Synuclein Accelerate Amyloid Formation. ACS Chem Neurosci 2022, 13 (1), 143–150. DOI: 10.1021/acschemneuro.1c00666.

5 Mesias, V. S. D.; Zhu, H.; Tang, X.; Dai, X.; Liu, W.; Guo, Y.; Huang, J. Moderate Binding between Two SARS-CoV-2 Protein Segments and α-Synuclein Alters Its Toxic Oligomerization Propensity Differently. J. Phys. Chem. Lett. 2022, 13 (45), 10642–10648. DOI: 10.1021/acs.jpclett.2c02278.

6 Nystrom, S.; Hammarstrom, P. Amyloidogenesis of SARS-CoV-2 Spike Protein. J Am Chem Soc 2022, 144 (20), 8945–8950. DOI: 10.1021/jacs.2c03925.

7 Jana, A. K.; Lander, C. W.; Chesney, A. D.; Hansmann, U. H. E. Effect of an Amyloidogenic SARS-COV-2 Protein Fragment on α-Synuclein Monomers and Fibrils. J. Phys. Chem. B. 2022, 126 (20), 3648–3658. DOI: 10.1021/acs.jpcb.2c01254.

8 Chesney, A. D.; Maiti, B.; Hansmann, U. H. E. SARS-COV-2 spike protein fragment eases amyloidogenesis of α-synuclein. J. Chem. Phys. 2023, 159 (1). DOI: 10.1063/5.0157331.

9 Li, B.; Ge, P.; Murray, K. A.; Sheth, P.; Zhang, M.; Nair, G.; Sawaya, M. R.; Shin, W. S.; Boyer, D. R.; Ye, S.;, et al. Cryo-EM of full-length alpha-synuclein reveals fibril polymorphs with a common structural kernel. Nat. Commun. 2018, 9 (1), 3609. DOI: 10.1038/s41467-018-05971-2.

10 Frishman, D.; Argos, P. Knowledge-based protein secondary structure assignment. Proteins 1995, 23 (4), 566–579. DOI: 10.1002/prot.340230412.

11 van Zundert, G. C. P.; Rodrigues, J.; Trellet, M.; Schmitz, C.; Kastritis, P. L.; Karaca, E.; Melquiond, A. S. J.; van Dijk, M.; de Vries, S. J.; Bonvin, A. The HADDOCK2.2 Web Server: User-Friendly Integrative Modeling of Biomolecular Complexes. J Mol Biol 2016, 428 (4), 720–725. DOI: 10.1016/j.jmb.2015.09.014.

12 Honorato, R. V.; Koukos, P. I.; Jimenez-Garcia, B.; Tsaregorodtsev, A.; Verlato, M.; Giachetti, A.; Rosato, A.; Bonvin, A. Structural Biology in the Clouds: The WeNMR-EOSC Ecosystem. Front Mol Biosci 2021, 8, 729513. DOI: 10.3389/fmolb.2021.729513.

13 Abraham, M. J.; Murtola, T.; Schulz, R.; Páll, S.; Smith, J. C.; Hess, B.; Lindahl, E. GROMACS: High performance molecular simulations through multi-level parallelism from laptops to supercomputers. SoftwareX 2015, 1-2, 19–25. DOI: 10.1016/j.softx.2015.06.001.

14 Huang, J.; Rauscher, S.; Nawrocki, G.; Ran, T.; Feig, M.; de Groot, B. L.; Grubmuller, H.; MacKerell, A. D., Jr. CHARMM36m: an improved force field for folded and intrinsically disordered proteins. Nat Methods 2017, 14 (1), 71–73. DOI: 10.1038/nmeth.4067.

15 Jorgensen, W. L.; Chandrasekhar, J.; Madura, J. D.; Impey, R. W.; Klein, M. L. Comparison of simple potential functions for simulating liquid water. The Journal of Chemical Physics 1983, 79 (2), 926–935. DOI: 10.1063/1.445869.

16 Bussi, G.; Donadio, D.; Parrinello, M. Canonical sampling through velocity rescaling. J Chem Phys 2007, 126 (1), 014101. DOI: 10.1063/1.2408420.

17 Parrinello, M.; Rahman, A. Polymorphic transitions in single crystals: A new molecular dynamics method. Journal of Applied Physics 1981, 52 (12), 7182–7190. DOI: 10.1063/1.328693.

18 Miyamoto, S.; Kollman, P. A. Settle: An analytical version of the SHAKE and RATTLE algorithm for rigid water models. Journal of Computational Chemistry 2004, 13 (8), 952–962. DOI: 10.1002/jcc.540130805.

19 Hess, B.; Bekker, H.; Berendsen, H. J. C.; Fraaije, J. G. E. M. LINCS: A linear constraint solver for molecular simulations. Journal of Computational Chemistry 1997, 18 (12), 1463–1472. DOI: 10.1002/(sici)1096-987x(199709)18:12<1463::Aid-jcc4>3.0.Co;2-h.

20 Humphrey, W.; Dalke, A.; Schulten, K. VMD: Visual Molecular Dynamics. J. Mol. Graph. 1996, 14 (1), 33–38. DOI: 10.1016/0263-7855(96)00018-5.

21 Bellaiche, M. M. J.; Best, R. B. Molecular Determinants of Aβ_42_ Adsorption to Amyloid Fibril Surfaces. J. Phys. Chem. Lett. 2018, 9 (22), 6437–6443. DOI: 10.1021/acs.jpclett.8b02375.

