## Supplementary material for "The Effect of SARS-COV-2 Protein Fragments on the Dimerization of α-Synuclein": Folder with atomic coordinates of start and final configurations of all trajectories in PDB format: README.txt

This directory contains (besides this README.txt) three folders (Rod Binding Dimer, Twister Binding Dimer, and Beta Strand Dimer) that contain files with the atomic coordinates (in PDB format) of aS structures discussed in the article. An additional folder (Parameter Files) contains mdp files of parameter settings for GROMACS simulations. 

Rod Binding Dimer;
This folder contains three sub-directories containing all Rod Binding dimer configurations:

1) Control:
Atomic coordinates (in PDB format) of the start and final configurations of the control dimer simulations:
r-c-1-initial : Initial structure of aS Rod Binding dimer replica 1
r-c-2-initial : Initial structure of aS Rod Binding dimer replica 2
r-c-1-final   : Final structure of aS Rod Binding dimer replica 1
r-c-2-final   : Final structure of aS Rod Binding dimer replica 2

2) with FI10:
Atomic coordinates (in PDB format) of the start and final configurations of the dimer simulations with FI10:
r-FI10-1-initial : Initial structure of aS Rod Binding dimer with FI10 replica 1
r-FI10-2-initial : Initial structure of aS Rod Binding dimer with FI10 replica 2
r-FI10-1-final   : Final structure of aS Rod Binding dimer with FI10 replica 1
r-FI10-2-final   : Final structure of aS Rod Binding dimer with FI10 replica 2

3) with SK9:
Atomic coordinates (in PDB format) of the start and final configurations of the dimer simulations with SK9:
r-SK9-1-initial : Initial structure of aS Rod Binding dimer with SK9 replica 1
r-SK9-2-initial : Initial structure of aS Rod Binding dimer with SK9 replica 2
r-SK9-1-final   : Final structure of aS Rod Binding dimer with SK9 replica 1
r-SK9-2-final   : Final structure of aS Rod Binding dimer with SK9 replica 2


Twister Binding Dimer;
This folder contains three sub-directories containing all Twister Binding dimer configurations:

1) Control:
Atomic coordinates (in PDB format) of the start and final configurations of the control dimer simulations:
t-c-1-initial : Initial structure of aS Twister Binding dimer replica 1
t-c-2-initial : Initial structure of aS Twister Binding dimer replica 2
t-c-1-final   : Final structure of aS Twister Binding dimer replica 1
t-c-2-final   : Final structure of aS Twister Binding dimer replica 2

2) with FI10:
Atomic coordinates (in PDB format) of the start and final configurations of the dimer simulations with FI10:
t-FI10-1-initial : Initial structure of aS Twister Binding dimer with FI10 replica 1
t-FI10-2-initial : Initial structure of aS Twister Binding dimer with FI10 replica 2
t-FI10-1-final   : Final structure of aS Twister Binding dimer with FI10 replica 1
t-FI10-2-final   : Final structure of aS Twister Binding dimer with FI10 replica 2

3) with SK9:
Atomic coordinates (in PDB format) of the start and final configurations of the dimer simulations with SK9:
t-SK9-1-initial : Initial structure of aS Twister Binding dimer with SK9 replica 1
t-SK9-2-initial : Initial structure of aS Twister Binding dimer with SK9 replica 2
t-SK9-1-final   : Final structure of aS Twister Binding dimer with SK9 replica 1
t-SK9-2-final   : Final structure of aS Twister Binding dimer with SK9 replica 2


Beta Strand Dimer;
This folder contains three sub-directories containing all Beta Strand dimer configurations:

1) Control:
Atomic coordinates (in PDB format) of the start and final configurations of the control dimer simulations:
b-c-1-initial : Initial structure of aS Beta Strand dimer replica 1
b-c-2-initial : Initial structure of aS Beta Strand dimer replica 2
b-c-1-final   : Final structure of aS Beta Strand dimer replica 1
b-c-2-final   : Final structure of aS Beta Strand dimer replica 2

2) with FI10:
Atomic coordinates (in PDB format) of the start and final configurations of the dimer simulations with FI10:
b-FI10-1-initial : Initial structure of aS Beta Strand dimer with FI10 replica 1
b-FI10-2-initial : Initial structure of aS Beta Strand dimer with FI10 replica 2
b-FI10-1-final   : Final structure of aS Beta Strand dimer with FI10 replica 1
b-FI10-2-final   : Final structure of aS Beta Strand dimer with FI10 replica 2

3) with SK9:
Atomic coordinates (in PDB format) of the start and final configurations of the dimer simulations with SK9:
b-SK9-1-initial : Initial structure of aS Beta Strand dimer with SK9 replica 1
b-SK9-2-initial : Initial structure of aS Beta Strand dimer with SK9 replica 2
b-SK9-1-final   : Final structure of aS Beta Strand dimer with SK9 replica 1
b-SK9-2-final   : Final structure of aS Beta Strand dimer with SK9 replica 2


Parameter Files;
This folder contains five mdp files with parameter settings:

ions.mdp          : Minimal parameter file for system preparation before ion addition
energy_min.mdp    : Energy minimization parameters
nvt-eqb.mdp       : Parameters for constant volume/temperature (NVT) equilibration
npt-eqb.mdp       : Paramters for constant pressure/temperature (NPT) equilibration
md-production.mdp : Molecular dynamics parameters for production simulation runs
